# Enhanced Transcriptome Maps from Multiple Mouse Tissues Reveal Evolutionary Constraint in Gene Expression for Thousands of Genes

**DOI:** 10.1101/010884

**Authors:** Dmitri D. Pervouchine, Sarah Djebali, Alessandra Breschi, Carrie A. Davis, Pablo Prieto Barja, Alex Dobin, Andrea Tanzer, Julien Lagarde, Chris Zaleski, Lei-Hoon See, Meagan Fastuca, Jorg Drenkow, Huaien Wang, Giovanni Bussotti, Baikang Pei, Suganthi Balasubramanian, Jean Monlong, Arif Harmanci, Mark Gerstein, Michael A. Beer, Cedric Notredame, Roderic Guigó, Thomas R. Gingeras

## Abstract

We characterized by RNA-seq the transcriptional profiles of a large and heterogeneous collection of mouse tissues, augmenting the mouse transcriptome with thousands of novel transcript candidates. Comparison with transcriptome profiles obtained in human cell lines reveals substantial conservation of transcriptional programs, and uncovers a distinct class of genes with levels of expression across cell types and species, that have been constrained early in vertebrate evolution. This core set of genes capture a substantial and constant fraction of the transcriptional output of mammalian cells, and participates in basic functional and structural housekeeping processes common to all cell types. Perturbation of these constrained genes is associated with significant phenotypes including embryonic lethality and cancer. Evolutionary constraint in gene expression levels is not reflected in the conservation of the genomic sequences, but is associated with strong and conserved epigenetic marking, as well as to a characteristic post-transcriptional regulatory program in which sub-cellular localization and alternative splicing play comparatively large roles.

---

Approximately ninety million years of evolution separate the mouse and the human genomes. During this period, selected and neutral genetic changes have accumulated, resulting in 60% nucleotide divergence^1^. Structural and coding organization, however, have been substantially maintained with approximately 90% of the mouse and human genomes partitioning into regions of conserved synteny, and more than 15,000 protein-coding orthologs (about 80% of all protein-coding genes) shared between these two genomes^2, 3^. Substantial information on the functional elements encoded in the human genome has been accumulated over the years. However, despite considerable effort^4, 5^, the mouse genome remains, in comparison, poorly annotated.

Here we characterized the transcriptional profiles from a diverse and heterogeneous collection of fetal and adult mouse tissues by RNA-seq. Using this data in conjunction with other data recently published^6^, we extended the mouse gene and transcript candidate set, and enhanced the current set of orthologous genes between these genomes to include long noncoding RNAs (lncRNAs) and pseudogenes. We also compared the mouse expression profiles with expression profiles in human cell lines, obtained in the framework of the ENCODE project, using identical sequencing and analysis protocols^7, 8^. While the compared profiles do not correspond to matched biological conditions, preventing the investigation of the evolutionary conservation of cell type versus species-specific transcriptional patterns, they allow for an investigation of the conservation of transcriptional features that are independent of the cell types specifically monitored. In particular, we have identified a well-defined subset of genes, the expression of which remains relatively constant across the disparate mouse tissues and human cell lines investigated here. Comparison with transcriptional profiles in multiple tissues of other vertebrate species^9, 10^ reveals that the constraint in expression has likely been established early in vertebrate evolution. Genes with constrained expression capture a relatively large and constant proportion of the RNA output of differentiated cells but not of undifferentiated cells, and is the main driver of the notable conservation of transcriptional profiles reported between human and mouse^2, 11, 12^ and other mammals^13^. Our analysis further shows that these genes are under specific conserved transcriptional and post-transcriptional regulatory programs.

## Expanded mouse transcriptional annotations

A total of 30 mouse embryonic and adult tissue samples and 18 human cell lines (generated as part of the human ENCODE project^7^) were used as sources for the isolation of polyadenylated (poly A+) long (>200 nucleotides [nt]) RNAs (Table S1), which were sequenced in two biological replicates to an average depth of 450 million reads per sample. Sequence reads were mapped and post-processed to quantify annotated elements in GENCODE^14^ (human v10, hg19) and ENSEMBL^15^ (mouse ens65, mm9), and to produce *de novo* transcriptional elements as previously described^8^. Reproducibility between replicates was assessed using a non-parametric version of the Irreproducible Discovery Rate (IDR) statistical test^8^ (Supplementary information, Tables S2 and S3A,B).

Reflecting the less developed state of the annotation of the mouse genome, GENCODE (v10) includes 164,174 long human transcripts, compared to 90,100 long mouse transcripts included in ENSEMBL (v65). By combining transcript predictions obtained using Cufflinks^16^ in our sequenced RNA samples with CAGE tag clusters recently produced by the FANTOM project^6^, we have identified about 150,000 novel transcripts in human^8^, and 200,000 in mouse (Table S3B), leading to similar numbers of transcripts in the two species, as illustrated by a few examples in Figure 1 (Supplementary information, Table S3C). Additionally, the mapping of the novel mouse transcripts back to the human genome lead to the discovery of 38 novel human genes not included in the models derived from human RNA-seq data, but supported by CAGE clusters. This underlines the importance of comparative approaches in completing genome annotations. By directly using the split RNA-seq reads at a stringent entropy threshold (Supplementary Information), we identified a set of about 400,000 highly confident splice junctions in the mouse genome, of which about half are novel. In contrast to annotated junctions, novel junctions are highly tissue-specific (Figure S1, Table S4A). By comparing to splice junctions in human, and using one-to-one whole genome maps^17^, we have assembled a set of 204,887 orthologous splice junctions (Table S4B). Moreover, we combined genome annotations and RNA-seq evidence and identified 3,641 mouse genes with antisense transcription (Supplementary Information)—a proportion of which, larger than expected by chance, are orthologous to human genes with antisense transcription (p-val < 10^-16^) (Table S5), indicating conservation in the occurrence of antisense transcription at human mouse orthologous loci^18^ (Figure 1B). Examples in Figure S2 and Table S6 show potential novel cases of a regulatory mechanism recently described involving antisense lncRNAs that contain a SINE element^19^.

**Figure 1.**
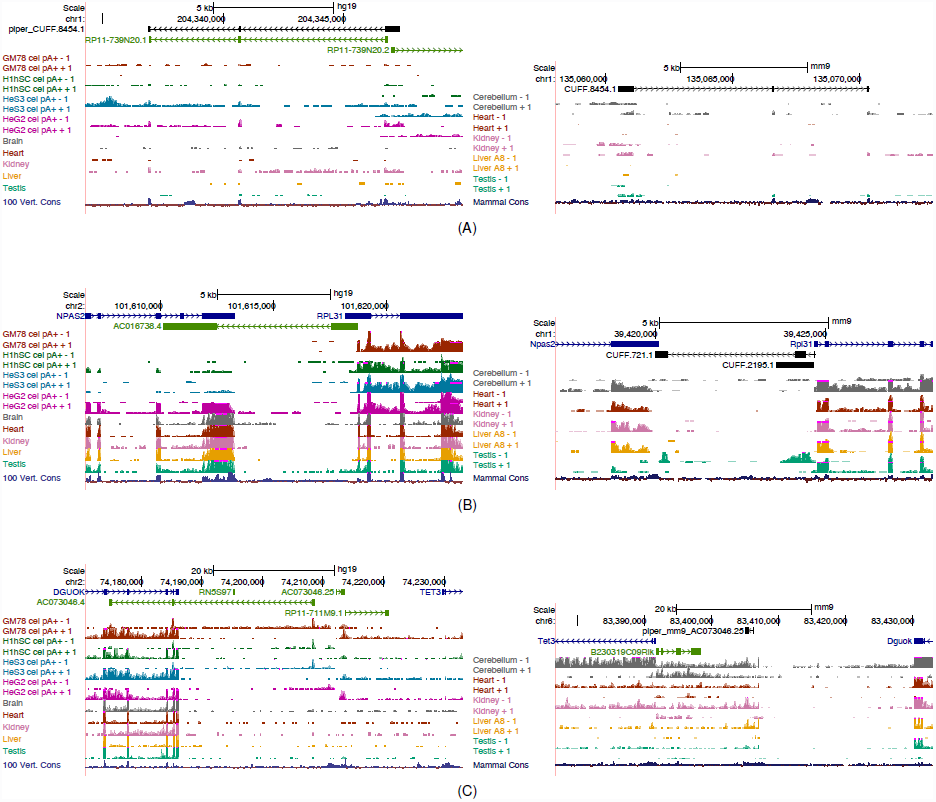
Exemples of extensions to the mouse annotation (left: human; right: mouse). **(A)** The novel intergenic transcript model CUFF.8454.1 has been inferred from mouse RNA-seq data in a region of the mouse genome without gene annotations. Mapping of this transcript model to the human genome reveals that it is the homologue of the lncRNA RP11-739N20.1 annotated in the human genome. **(B)** Two transcript models, CUFF.721.1 and CUFF.2195.1 have predicted antisense to two neighboring protein-coding loci (Npas2 and RPL31). RNA-seq shows that their expression is restricted to testis. CUFF.721.1 is the likely homologue of the annotated human AC016738.4, which is antisense and overlaps the human ortholog of Npas2 and RPL31. In human, AC016738.4 appears to be also expressed specifically in testis, although this is not conclusive, since we lack stranded RNA-seq data. **(C)** The mapping of the human AC0703046.25 lncRNA to the mouse genome identified a potential mouse transcript in the syntenic region between the DGU0K and the TET3 genes (piper_mm9_AC073046.25). This transcript was not included in our predicted models, but it has strong support by RNA-seq data, specifically in cerebellum, kidney and liver. Tissue RNA-seq data in human is from Ilumina Body Map (HBM, Refs). Plots are UCSC browser screenshots where novel mouse models are indicated in black, human Gencode annotation in blue, green and red, and mouse CSHL and human HBM RNA-seq signal in different colors at the bottom. Annotated genes are represented by the longest transcript.

Using one-to-one whole-genome maps and sequence homology, we identified a set of 851 human-mouse orthologous long non coding RNA (lncRNA) genes (Figure 1, Supplementary Data Files and Information). Of these, only 189 overlap with the set of 2,736 one-to-one human-mouse lncRNAs recently described^20^, reflecting the yet incomplete characterization of mammalian lncRNAs, and of lncRNA orthology. Using localization data in human cell lines^8^ we found these genes to segregate in two clearly distinct nuclear vs. cytosolic populations (Figure S3A). Expression levels of orthologous lncRNAs correlate weakly with phylogenetic depth (i.e. the number of mammalian species in which a given lncRNA can be detected, Figure S3B). While lncRNAs show distinct tissue- and species-specific expression patterns, we identified twelve lncRNAs expressed in at least 50% of the samples in each species. This small set of conserved broadly expressed genes may play important functions in mammalian cells (see Figure 1C for an example). We found these genes to be highly enriched among nuclear lncRNAs (Figure S3A). Additionally, we identified a set of 129 orthologous pseudogenes, of which 32% are expressed in one species and 4% in both (Supplementary Information).

## Genome-wide conservation of gene expression and splicing

There is overall, substantial genome-wide conservation of expression levels between human and mouse irrespective of the cell or tissue type of the sample. We computed genomewide expression profiles, measured as average read density, for all orthologous 100-nt bins spaced equally along the human and mouse genomes (Figure S4A,B, Supplementary Information). We found substantial correlation in average read density at orthologous bins (cc=0.67, Figure 2A). This correlation is significant not only for exonic regions^2^ (Figure S4C), but also for alignable intronic (Figure S4D) and intergenic regions (Figure 2B). However, most of this intergenic transcription is proximal to annotated genes (41% less than 10kb from the closest annotated gene termini, Figure 2C). This is partially the consequence of the decreasing number of intergenic bins with distance to the closest gene (Figure S5). In any case, the murine-human expression correlation decays with distance to the closest annotated gene (Figure 2D). Permissive transcription close to protein coding genes could be the origin of many lncRNAs. However, when computing the distance between annotated lncRNAs and the closest neighbor annotated gene we found this distance to be larger than for protein coding genes (means of approximately 66Kb and 35Kb respectively). Expression levels correlate with phylogenetic conservation, as measured by phastCons^21^ scores^22^ (Figure 2E). However, a fraction of orthologous bins having low sequence conservation are still densely transcribed (5% of the least conserved bins have read density greater than 10) and the bins that correspond to higher expression include a wide range of sequence conservation values (Figure 2F). Highly expressed intergenic bins are slightly enriched for GWAS hits (p-val≈0.055), and strongly enriched for cis-eQTLs (p-val<2.2e-16, Supplementary Information), the latter suggesting an important role for enhancer transcription in the regulation of gene expression. There is also substantial conservation of antisense transcription^18^. For each sense/antisense orthologous gene pair we computed the ratio of antisense-to-total gene expression averaged over all conditions, and found strong correlation of average antisense-to-total gene expression ratio in orthologous genes (cc=0.68) as well of its variation among samples (cc=0.52) (Figures 3A and S6). Antisense transcription has been suggested as an important regulatory mechanism^23^, and our results indicate that it may have been conserved over large evolutionary distances.

**Figure 2.**
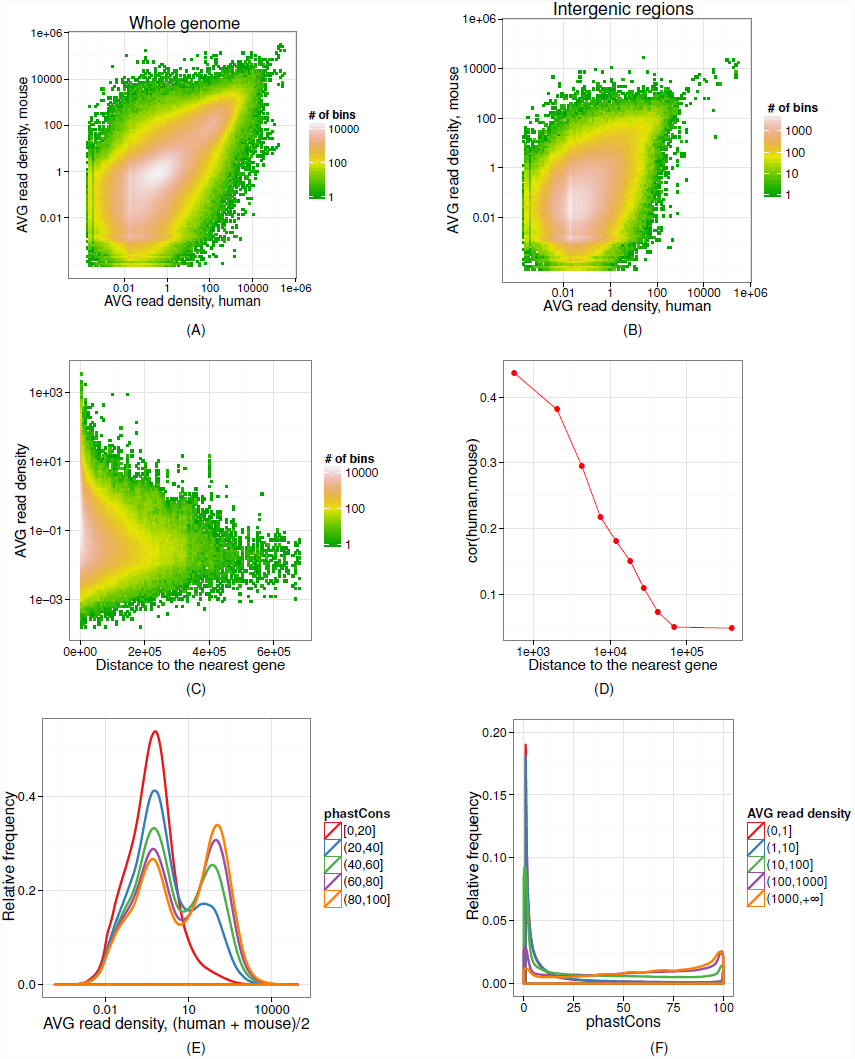
Genome wide conservation of expression profile. **(A)** The joint distribution of log_10_ average read density in orthologous 100-nt bins in human (x-axis) and in mouse (y-axis); cc=0.67. **(B)** The distribution in (A) limited to intergenic regions; cc=0.37. **(C)** The distribution of log_10_ average read density in intergenic regions (average between human and mouse) as a function of distance from the nearest gene. **(D)** Correlation of log_10_ average read density in human and mouse as a function of distance from the nearest gene. **(E)** The distribution of log_10_ average read density in 100-nt bins as a function of phastCons score (conservation score across 45 vertebrate species). **(F)** The distribution of phastCons score of a 100-nt bin as a function of the average read density.

Conservation of the exonic structure of human-mouse orthologous genes, as well as of the splice site sequences has also been reported^24^. RNA-seq further allows the investigation of the patterns of usage of exons and splice junctions. We computed the average inclusion, measured as percent-spliced-in (PSI, ψ), and found strong correlation of inclusion at orthologous splice junctions (cc=0.59, Figure 3B). The correlation is mostly driven by relatively low average inclusion levels of junctions annotated as alternative in both human and mouse. These junctions constitute a large part of the orthologous set (45%) and are used more variably in both species (Figure 3C). In addition, we computed average splicing processivity, measured using the completeness of splicing index (coSI, θ^25, 26^), and found also significant correlation, albeit lower (cc=0.35, Figure S7A,B). This is expected, since the inclusion level of an exon in the final RNA product is likely to be physiologically more relevant than the efficiency with which the exon is included.

**Figure 3.**
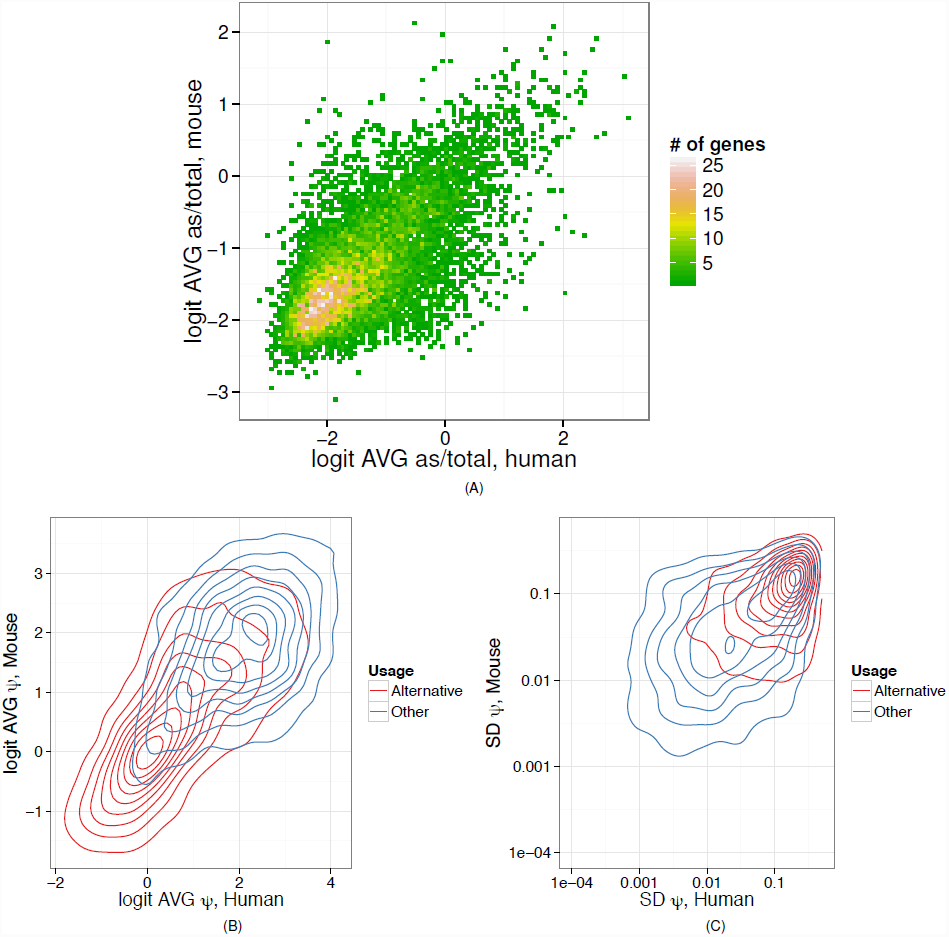
**(A)** The joint distribution of the average antisense-to-total expression ratio (the number of reads mapped to the opposite strand as a fraction of the number of reads mapped to both strands) in pairs of orthologous protein-coding genes; cc=0.68. **(B and C)** Contour plots of the joint probability distribution of the average usage (ψ, percent-spliced-in) of Splice Junctions (SJs) (B), and of the standard deviation of SJ usage (C) in orthologous SJs pairs. Logistic transformation *(logit)* was used in (B). SJ with constant complete inclusion or exclusion are not shown. “Alternative” denotes SJs that are annotated alternative in both species.

## Evolutionary conserved constraint in gene expression

As previously reported^8, 27^, we found that within a given cell population the levels ofgene expression may vary up to six orders of magnitude (Figure S8). In contrast however, the expression of a given gene across cell types varies relatively little. Using a two-factor ANOVA, with gene and cell type as two factors, we found that the variance across genes accounts for 76% and 71% of the total variance of gene expression across human and mouse cell types, respectively, while the fraction of the variance that can be directly attributed to cell type is less than 1% (Supplementary Information). Indeed, we found a large fraction of genes, the expression of which varies relatively little across tissues and species. In Figure 4A we computed the distributionof the dynamic range of gene expression (DNR) in orthologous genes across the entire set of human and mouse samples. For each gene expressed in at least two samples both in humanand mouse, we computed the log_10_ ratio between the highest and lowest measured expression. The distribution is bimodal, uncovering two broad gene classes. This is not an effect of comparing mouse tissues with human cell lines, since the same pattern is obtained when using RNA-seq obtained in human tissues by the Illumina Body Map project (Figure S9). We decompose the distribution assuming two underlying Gaussians, and we took DNR=2 at the approximate intersection point (Figure S10, Supplementary Information). In this way, we obtained a set of 6,636 genes (∼40% of all 15,736 orthologous genes, Figure 4B), the expression of which remains relatively constant (i.e. within two orders of magnitude) across species and cell types. Genes with constrained expression show wider expression breadth (Figure S11A) and less tissue specificity than the rest of the orthologs (less than 10% of tissue specific orthologs are included in this set, Supplementary Information). However, they can be eventually detected as differentially expressed at a rate similar to the rest of the orthologs (82% vs. 89%). They also show higher expression levels (Figure S11B). Therefore, although they representonly about 17% of all annotated genes, they capture a high proportion of the poly A+ transcription in the cell (39% on average in human and 41% in mouse), a proportion that remains remarkably constant across all tested human and mouse cell types (Figure 4C). Mouse embryonic cell lines are an exception, constrainedgenes generating only about 20% of the cell’s transcriptional output. We also founda negative association between minimal expression and DNR (Figure S12). Thus, genes with constrained expression tend to have also higher minimal expression, suggesting that thesegenes in their default state may already be primed for transcription. To eliminate gene expression as a potential confounding factor, most downstream analyses have been carried out in a subset of 5,519 genes with constrained expression, and in a subset of identical size from the rest of the orthologous genes for which we could compute DNR, with matched expression in human and mouse (the unconstrained set, Figure S11C, Supplementary Information).

**Figure 4.**
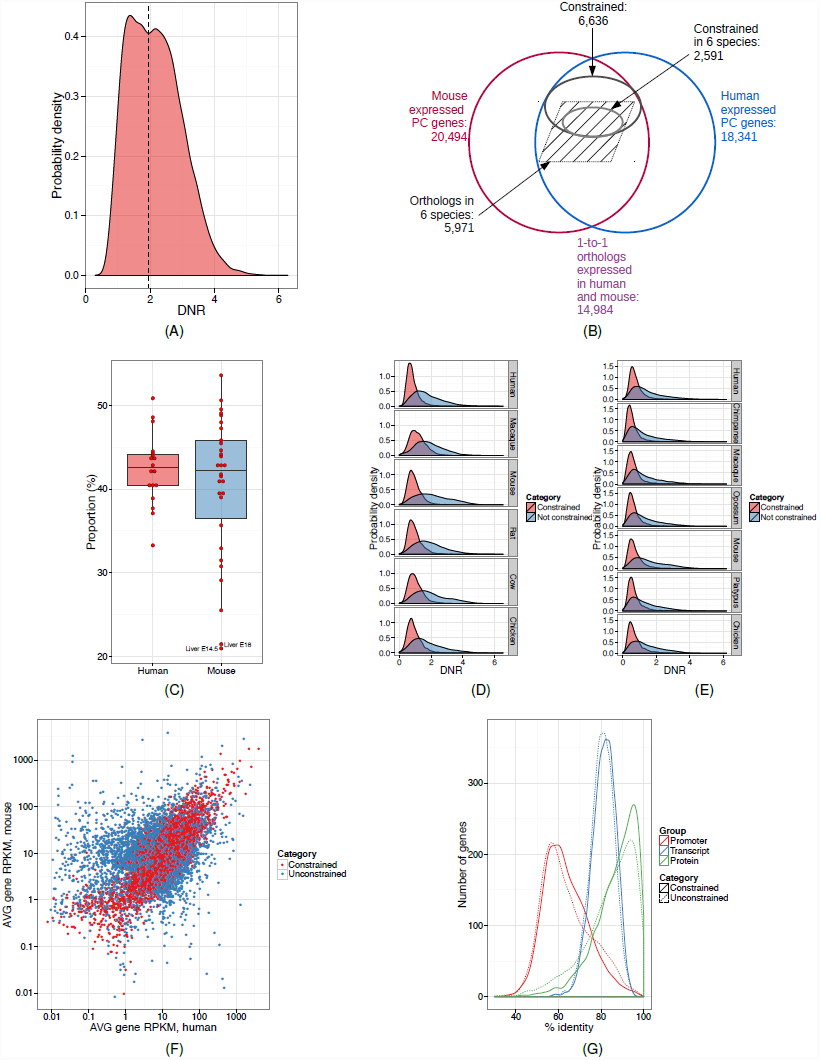
**(A)** The distribution of the dynamic range (DNR, log_10_ of the ratio of the largest and the lowest non-zero observation) of gene expression level in orthologous genes across human and mouse samples. **(B)** Venn diagram of the relationship between orthologous and constrained genes. **(C)** Proportion of nucleotides in expressed genes, as assessed by PolyA+ RNA-seq, that originates from constrained genes in human cell lines and mouse tissues. The labeled outliers correspond to mouse embryonic samples. **(D and E)** The distribution of DNR in human/mouse constrained and unconstrained genes in Merkin et al.^10^ (D) and Barbosa-Morais et al.^9^ (E). **(F)** The joint distribution of log_10_ average gene RPKM in pairs of orthologous protein-coding genes; constrained genes are shown in red. **(G)** The distribution of promoter, transcript, and protein pairwise sequence identity between human and mouse in constrained and unconstrained genes.

Using transcriptome data recently published in multiple tissues across a number of vertebrate species^9, 10^, we found that the expression of these genes is actually constrained beyond human and mouse, and seems to have been established early in vertebrate evolution. About 94-98% of genes with constrained expression in human and mouse, have DNR across tissues less than two in other vertebrate species (Figure 4D, E). We further identified a collection of 2,591 genes whose expression levels show relatively little variation across different cell types and species within vertebrates (out of the 5,971 orthologous across these species that are expressed in both mouse and human^10^ (Figure 4B). Gene Ontology analysis reveals that genes with constrained expression participate in basic functional and architectural housekeeping processes common to all cell types (Supplementary data archive 5). These genes are the main drivers of the substantial conservation of expression reportedinorthologous human and mouse genes^28-30^. Indeed, the correlation of average gene expression between human and mouse measured in the set of constrained genes is 0.82, compared to only 0.32 for the unconstrained gene set (Figure 4F).

Constraint of gene expression is not reflected in sequence conservation of either the gene body or the proximal promoter regions^31^ (p-val ≈0.11 for the difference in phastCons scores; Figure 4G). In contrast, we found that constrained genes exhibit characteristic patterns of histone modifications, quite divergent from that of unconstrained genes. Using the data collected in these and the human ENCODE studies^3, 7^ (Table S7A, B, Supplementary Information), we computed genic profiles of normalized histone modification signals, averaging them over all studied cell types (Figure 5A). We found much stronger signal for active histone modifications (H3K4me3, H3K27ac, and H3K36me3) in constrained compared to unconstrained genes. Since the set of genes in which we performed the analysis are controlled for differences in gene expression, thisis not the result of constrained genes exhibiting higher expression levels. Furthermore,we controlled by gene expression within each sample separately, and found the same effect(Figure S13). While the association between gene expression and chromatin structure is well known, and models have been developed able to predict gene expression from levels of histone modifications with high accuracy^32, 33^, our results suggest that strong chromatinmarking is not only associated to high expression levels, but also to high transcription stability across cell types and tissues. This association has been apparently conserved during evolution, since we found that the conservation of the active epigenetic signals is stronger in constrained compared to unconstrained genes (Figure 5B). These results suggest that constrained and unconstrained genes may be under globally different epigenetic programs. Further supporting this, we found that constrained genes are regulated by broad promoters (as defined by the FANTOM consortium^6, 34^) more often than unconstrained genes (67% vs 52%, p-val ≈0), that their promoters host more transcription factor (TF) ChIP-seq peaks (using data collected in the ENCODE project^7^, Supplementary Information), than unconstrained genes (Figure 5C), and are slightly depleted in repeat elements at their TSS (density of repeat elements in the promoter region 0.82 vs. 0.87, p-val ≈0.03, Supplementary Information). Moreover, when performing principal component analysis to classify the promoters of human mouse orthologous genes based on ChIP-seq measured binding strength of TFs, we found a separation between promoters of constrained and unconstrained genes, although the components capture only a small fraction of the whole variance (Figure 5D).

**Figure 5.**
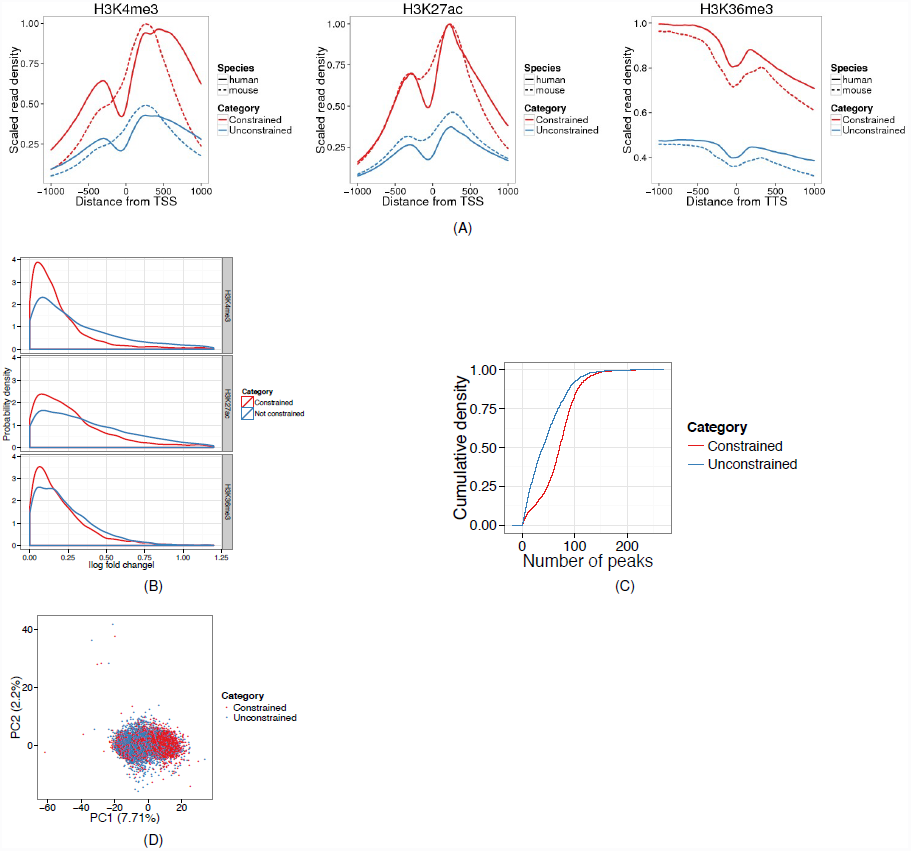
**(A)** Normalized histone modification profiles 1Kb around Transcription Start Site (TSS) for H3K4me3 and H3K27ac, and around the Transcription Termination Site (TTS) for H3K36me3 in constrained and unconstrained genes. **(B)** The difference in normalized average histone modification signals 1Kb around TSS (|Δlog Signal|) in constrained and unconstrained genes **(C)** The cumulative distribution of the number of TF ChIP-seq peaks in promoter regions of human constrained and unconstrained genes. **(D)** Principal component analysis of ChIP-seq measured binding strength of TFs in constrained and unconstrained genes.

The maintenance of constrained RNA production levels appears to have also impacted post-transcriptional regulation. First, genes with constrained gene expression tend to be constrained also in their cellular localization. That is, their ratio of cytosolic versus nuclear abundance across human cell lines is much less variable that for unconstrained genes (Figure 6A). Moreover, transcripts from constrained genes tend to be enriched in the cytosol compared to transcripts from unconstrained genes (70% of the constrained genes are mostly found in cytosol compared to 60% of unconstrained ones (p-val≈0, Supplementary Information). Second, we found that splicing plays a comparatively more important role determining the cellular abundance of transcript isoforms in constrained than in unconstrained genes. Gonzalez-Porta et al.^35^ developed a method to estimate the fraction of the variance in transcript abundance that can be explained by variance in gene expression, when measured across a number of samples (cell types). Here we found that this fraction is on average 0.81 in human cell lines, and 0.82 in mouse tissues—indicating that overall regulation of gene expression plays the major role in defining cell type specificity. This is consistent with tissue dominated clustering of gene expression profiles compared with the species dominated clustering of splicing profiles^9, 10^. We have found, however, substantial differences between constrained and unconstrained genes. The average fraction of the variance in transcript abundance explained by gene expression is 0.76 and 0.79for constrained genes, in human and mouse respectively, while these values are 0.91 for both species for unconstrained genes (p-val≈0, Figure 6B). Thus, regulation through splicing appears to play comparatively a more important role in constrained than unconstrained genes.

**Figure 6.**
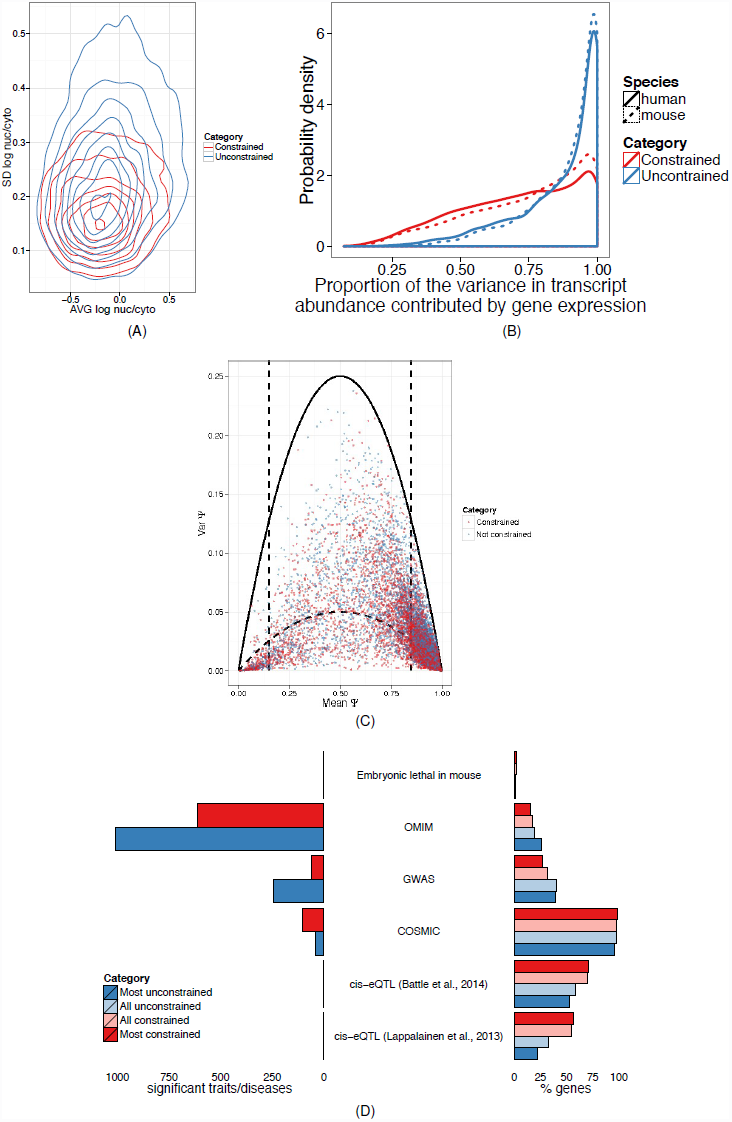
**(A)** Contour plots of the joint probability distribution of the average vs. standard deviation (SD) of log_10_ nuclear-to-cytosolic ratio in constrained and unconstrained genes. **(B)** Distribution of the proportion of the variance in transcript abundance across human and mouse samples that can be explained by the overall variance in gene expression. Values close to 1 indicate that changes in abundance of transcript isoforms originate mostly from changes in gene expression, values closeto 0 suggest that most of these changes are due to splicing changes in the relative proportion of transcript isoforms. (**C)** Mean versus Variance of splice junction inclusion (ψ) in constrained and unconstrained genes. To identify junctions with constrained inclusion at intermediate levels we set a threshold (inner parabola) of 20% of the maximum variance for the given mean in a Bernoulli distribution (outer parabola) in the interval of mean inclusion [0.15, 0.85]. **(D)** The percent of all and of the1000 most constrained and unconstrained genes (right) that are lethal in mice accordingtothe Jax mice embryonic lethality database^48^, have hits in OMIM^44^, the NHGRIGWAS catalog^45^, COSMIC^43^, and that have eQTLs^46, 47^. On the left, the numberofsignificant traits/diseases associated to constrained and unconstrained genes in OMIM,theNHGRI GWAS catalog and COSMIC.

Consistent with the relatively minor role of splicing in defining cell type specificity, compared to expression, we found that about 83% or orthologous splice junctions (65,485 out of 78,602 in our set of constrained and unconstrained genes matched by expression) are systematically included at inclusion levels ψ larger than 0.85 in all human and mouse samples. We have also found a small set of 123 junctions with extremely low inclusion levels (ψ < 0.15) in all human and mouse samples. Among the remaining ∼20,000 junctions we used an approach parallel to that for gene expression, to identify junctions the inclusion of which is constrained across cell types and species. By setting a threshold on the variance of ψ to be 20% of the maximum possible variancefor a Bernoulli distribution with the given mean, we have identified a set of 1,430 orthologous splice junctions with inclusion constrained at intermediate levels (0.1 < ψ < 0.9, Figure 6C, S14) across all human and mouse samples. That is, these are junctions with similar inclusion levels in allhuman and mouse samples investigated here. A majority (59%) of these junctions constrained in splicing belong to genes that are also constrained in expression, consistent with the comparatively more important role of splicing in the regulation of these genes.

## Conclusion

“Housekeeping” genes are characterized as genes involved in the maintenance of a cell’s basic functioning and are expressed in a ubiquitous and uniform fashion in different biological conditions ^36^. Numerous collections of housekeeping genes have been proposed ^36-38^, but the membership overlap is only moderate (Table S8). Here, by probing gene expression simultaneously across biological conditions and species – and therefore by introducing an evolutionary component – we found a principled way to identify a set of genes with constrained gene expression. While not identical, this set of genes has considerable overlap with two of the largest housekeeping gene sets recently reported: 43% of our genes are included in the set of 3,664 housekeeping genes reported by Eisenberg andLevanon (E-L)^36^, and 69% in the set of6,560 stably expressed that we have derived from the data reported by FANTOM5 (F5)^6^ (Supplementary information and Figure S15).

Here we have found that genes with evolutionary constrained levels of expression account for the bulk of transcription in differentiated mammalian cells, meaning that a substantial fraction of the RNA content in mammalian cells remains remarkably constant across large biological and evolutionary scales. While the evolutionary forces responsible for this transcriptional stability have not left an obvious imprint in the sequence of mammalian genomes^29^, we have found, in contrast, a remarkable epigenetic signature. Our results show that the evolutionary stability in gene expression is associated with strong and consistent marking by active histone modifications. This indicates that chromatin marking is not only a reflection of transcriptional levels, but also of transcriptional stability. This marking, moreover, is conserved across the evolutionary distance separating human and mouse. Genes with constrained expression seem to have evolved, in addition, a characteristic program of post-transcriptional regulation, in which sub-cellular localization and alternative splicing play a more prominent regulatory role than in genes with unconstrained expression. These results are consistent with an evolutionary interplay between transcriptional regulation and regulation by splicing, in which the maintenance of tight expression levels would enhance the role of splicing as a mechanism to modulate the abundance of individual transcript isoforms. They also suggest that lack of sequence constraint (which is detectable over less than 10% of the mammalian genomes, according to recent estimates^39^) cannot be naively equated to lack of biological function^40^.

The essential role of genes with constrained expression in basic cellular architectureand function suggests that disruption of their functions is likely to have dramatic phenotypic consequences, including cell survival. While the relationship between housekeepinggenes and disease remains controversial^41, 42^, we indeed found that mutations in genes with constrained levels of expression are most often associated with embryonic lethality, and that somatic mutations in these genes are associated to a broad spectrum of cancerousphenotypes^43^ (Figure 6D). In contrast, mutations in unconstrained genes are more likely tobe associated with other diseases (as reported in OMIM^44^ and the NHGRI GWAS catalog^45^, Figure 6D), as expected, since mutationslethal during development are unlikely to cause detectable diseases. The larger numberof GWAS hits associated to unconstrained genes underscores the complexities involved in identifying disease causative genetic variations when these affect gene. We found that theexpression of unconstrained genes is also more variable across individuals compared to theexpression of constrained genes (Figure S16). Large expression variability decreases the power to identify expression QTLs (eQTLs), and indeed we observed that unconstrained genes, which are more often associated to diseases, are substantially depleted for eQTLs compared to genes with constrained levels of expression^46, 47^ (Figure 6D).

In summary, by introducing an evolutionary dimension, we identified in a principled way, a set of genes with constrained expression across cell types and species. These genes contribute to a core invariable component of mammalian transcriptomes, contributing thus little to cell type specificity and species differentiation. They are the main drivers ofmany transcriptional features that have been reported as conserved across mammalian transcriptomes.

## Acknowledgements

This work was supported by the National Human Genome Research Institute (NHGRI) grantsnumber 1U54HG007004-1, U41HG007000, U01HG004695, U54HG004555, and U41HG007234, by the Spanish Plan Nacional grants number BI02011-26205 and BFU2011-28575, the ERC grant number 294653, LaCaixa, and the EU-FP7 quantomics project. We would also like to thanks Fyodor Kondrashov for useful discussions, and Emilio Palumbo for technical support.

Author Contributions: T.R.G., R.G., and C.A.D. designed the project; T.R.G. R.G. and C.A.D. managed the project; L.-H.S prepared biological samples; L.-H. S. and M.F. preparedRNA samples; L-l.-H.S., J.D., and M.F. prepared Illumina RNA-seq libraries; C.Z. and J.L.administered computer infrastructure and quality control for data storage and analysis;D.D.P., S.D., A.B., A.D., P.P.B., G.B., B.P., S.B., J.M., A.T., A.H., M.A.B., C.N. analyzeddata; M.G., M.A.B. assisted with manuscript preparation, T.R.G., R.G., D.D.P., S.D., A.B.wrote the paper with input from all authors. All authors discussed the results and commented on the manuscript.

**Author Information** Sequences are available from the Short Read Archive and the Mouse ENCODE website, a list of accession numbers is given in Table S1.

The authors declare no competing financial interests.

The raw data (FASTQ), mapped data (BAM) and lists of quantified elements are availableat http://mouse.encodedcc.org/. These data, as well as additional data on all intermediate processing steps, are also available on the RNA Dashboard (http://genome.crg.cat/encode_RNA_dashboard/).

